# Anoikis resistance in mammary epithelial cells is mediated by semaphorin 7a Semaphorin-7A and anoikis resistance

**DOI:** 10.1101/2021.06.28.449786

**Authors:** Taylor R. Rutherford, Alan M Elder, Traci R. Lyons

## Abstract

Semaphorin-7a (SEMA7A), best known as a neuroimmune molecule, plays a diverse role in many cellular processes and pathologies. Here, we show that SEMA7A promotes anoikis resistance in cultured mammary epithelial cells through integrins and activation of pro-survival kinase AKT, which led us to investigate a role for SEMA7A during postpartum mammary gland involution—a normal developmental process where cells die by anoikis. Our results reveal that SEMA7A is expressed on live mammary epithelial cells during involution, that SEMA7A expression is primarily observed in α6-integrin expressing cells, and that luminal progenitor cells, specifically, are decreased in mammary glands of SEMA7A−/− mice during involution. We further identify a SEMA7A-α6/β1-integrin dependent mechanism of mammosphere formation and chemoresistance in mammary epithelial cells and suggest that this mechanism is relevant for recurrence in breast cancer patients.

## INTRODUCTION

It has been postulated that the functional plasticity of the adult mammary epithelium, which is required for response to hormonal changes during pregnancy and lactation, may make the breast tissue more susceptible to malignancy.Identifying mechanisms of normal mammary epithelial cell (MEC) biology that contribute to tumorigenesis is critically important for furthering our understanding of breast cancer and extending the survival of breast cancer patients. Of particular interest are postpartum breast cancers (PPBCs), defined as breast cancers diagnosed in women within 10 years of most recent childbirth, as these cancers are nearly three times more likely to result in lethal metastasis(1).

The majority of mammary morphogenesis occurs postnatally beginning at puberty and culminating with lactation following pregnancy. As such, differentiation of the mammary epithelium is not complete until a full pregnancy-lactation cycle. Upon weaning, or after pregnancy in the absence of nursing, the mammary gland regresses to resemble the pre-pregnant architecture through a process termed postpartum involution. In mice, postpartum involution occurs in two primary phases(2). In the first phase, milk proteins accumulate in the mammary ducts and alveoli resulting in downstream phosphorylation and activation of <u>s</u>ignal <u>t</u>ransducer and <u>a</u>ctivator of <u>t</u>ranscription (pSTAT3), and subsequent lysosomal-mediated cell death(3). In mice, if pups are returned within 48 hours of weaning, this process can be reversed, and lactation can continue. If pups are not returned, however, the irreversible second phase of involution begins. This phase is characterized by degradation of the mammary basement membrane by matrix metalloproteinases and anoikis-mediated cell death in the absence of laminin-mediated activation of integrin alpha-6/beta-1 (α6β1) and downstream pro-survival signaling(4–6).

SEMA7A, or CD108w, is a member of the semaphorin protein family, named after the English verb “semaphore”, which means to transmit a signal across a long distance. SEMA7A was first identified for its expression on lymphocytes(7), but has since been shown to exert diverse roles in neuronal development(8), cell differentiation(9–11), ECM deposition(12, 13), various components of the immune system(14–17), and pathologies including fibrosis(12, 18, 19) and cancer’(13, 16, 20–22). Though elevated SEMA7A is observed in breast cancer, where we have established a role for it in conferring cell survival, in part, via promotion of anoikis resistance(13, 23), the role of SEMA7A in normal MECs has not previously been investigated. SEMA7A is known to signal through β1-integrin in immune cells, neurons, and cancer; β1-integrin also mediates MEC survival during development, in part, via phosphorylation and activation of downstream pro-survival signaling mediated by PI3-kinase and protein kinase B, or Akt1(24). Though it is known that redundant secretory epithelial cells die during postpartum involution, the factor(s) that allow some cells to survive and regenerate a lactation-competent gland for future pregnancies remain unknown. Herein, we have identified enrichment for SEMA7A expressing α6β1 positive MECs during the matrix remodeling period of postpartum involution and a role for SEMA7A in specifically promoting the survival of α6-integrin+ luminal progenitor cells. We provide evidence that exogenous stimulation of immortalized MECs with purified SEMA7A protein promotes anoikis resistance via activation of α6 and β1-integrins and activation of phosphorylated AKT (pAKT). Consistent with a role for SEMA7A in promoting anoikis resistance, SEMA7A knockout mice (SEMA7A^tmalk1/J^) exhibit phenotypes associated with accelerated involution including early presence of apoptotic cell bodies in mammary lumens, increased pSTAT3, and premature reappearance of adipocytes. α6-integrin also drives maintenance of tissue resident stem cells and therapy resistant cancer stem cells(25) and our findings reveal that SEMA7A drives stem cell phenotypes and chemoresistance in cultured MECs in a manner that is dependent on α6 and β1. Finally, we provide evidence for SEMA7A in driving transformed phenotypes in normal MECs and reversing transformed phenotypes in genetically identical, ras-transformed, variants of the same cell line. Our results are the first to identify a role for SEMA7A in anoikis resistance and promotion of stem and cancer cell phenotypes via activation of α6 and β1-integrins.

## METHODS

### Cell culture

MCF10A/MCF12A cells were obtained from, and cultured, according to ATCC. MCF10DCIS.com cells were obtained from K. Polyak (University of Harvard, Cambridge, MA). Cells were validated by the DNA sequencing core at the University of Colorado Anschutz Medical Campus and found to be pure populations of their respective cell lines. Cells were regularly tested for mycoplasma throughout studies. Cells were sub-cultured as previously described (reference), or according to ATCC standards. Exogenous SEMA7A protein utilized in our studies was purified from media of cells stably transfected with an Fc-tagged version of SEMA7A by published methods(14, 26). Control plasmid (pcDNA3.1) was obtained from H. Ford (University of Colorado Anschutz Medical Campus, Denver, CO).

### shRNA Knockdown and Overexpression

shRNA plasmids targeting Sema7a, and a negative control shRNA (SABiosciences), were amplified in E. coli and plasmid DNA was isolated by Plasmid Maxi-Prep (Qiagen). MCF10DCIS.com-GFP cells were cultured overnight to approximately 80% confluence. 1 μg/μL of each shRNA was added to Transfectagro Reduced-Serum Medium (Corning) and incubated for 15 minutes with 4 μL of X-treme Gene HP DNA transfection reagent (Roche). Transfected cells were selected for by hygromycin resistance. Stable knockdown was confirmed by qPCR as shown in Black et al(27). Negative control cells were transfected with a scrambled artificial sequence not matching human, mouse, or rat. Overexpression plasmid (SEMA7A-Fc) was a generous gift from R. Medzhitov (Yale University, New Haven, CT). All other over-expression plasmids (p304-V5-Blasticidin and V5-SEMA7A)were obtained from the Functional Genomics Core at the CUAnschutz Medical Campus and overexpression was confirmed via qPCR and western blot analysis. KD (27) and overexpression (13) as previously described.

### Animal Studies

All animal procedures were approved by the University of Colorado Anschutz Medical Campus Institutional Animal Care and Use Committee. Sema7a^tm1Alk/J^ mice (a generous gift from Alex Kolodkin at John’s Hopkins University) and C57Bl/6 (Jax) were housed and bred as previously described.(28) Briefly, C57Bl/6 SEMA7A^tm1Alk^ females were bred with WT C57Bl/6 males to induce pregnancy, lactation, and postpartum involution. Pup numbers were normalized to 6-8 per dam after birth to ensure for adequate lactation. Postpartum involution was initiated by forced weaning of pups at the peak of lactation (L10-L12). #4 right and left mammary glands were harvested from age-matched nulliparous animals and postpartum dams at involution days 1,2,4,6,8,10,14, and 28. For flow cytometry, inguinal lymph nodes were removed prior to tissue digestion. #3 mammary glands were also harvested, formalin fixed, paraffin embedded, and 5 micron sections generated for histologic/immunohistochemical analaysis. See Supplemental Table 1 for animal study sample sizes.

### In vitro survival assays

MCF10A and MCF12A cells were cultured with 200ng/ml purified SEMA7A or PBS vehicle control in adherent (tissue culture plastic) or forced suspension conditions (ultra-low attachment plates (Corning, Corning, NY, USA: #3473)). Cells were seeded at a density of 1000 cells/well in a 96-well plate plus media only wells as controls. Cell death was measured 24 hours post seeding via luminescence using the Caspase Glo assay (Promega, Madison, WI, USA: #G8090). Annexin-V /7AAD staining (Biolegend, San Diego, CA, USA: #640930) and pAKTS473 (ThermoFisher: Waltham, MA, USA, #17-9715-42) were also used to confirm cell viability by flow cytometry. See flow cytometry methods for staining protocol. Function blocking antibodies for b1 (CD29-9EG7; BD Biosciences: #550531) and a6 integrin (ThermoFisher: #14-0495-82) were used to disrupt integrin signaling in the presence of SEMA7A or as antibody alone controls to determine off-target effects. IgG controls were also used. 9EG7 was used at a concentration of 0.6ug/ml while GoH3 was used at a concentration of 100ug/mL for inhibition studies consistent with previous reports (29, 30). Cells were cultured with anti-integrin inhibitors at time of seeding.

### Label retention stem cell assay

MCF10A and MCF12A cells were labeled with CellTrace Violet (ThermoFisher: #C34557) for 20 minutes at 37°C according to manufacturer’s instructions. A sample of labeled cells was analyzed by flow cytometry to determine labeling efficiency and fluorescent intensity at time of seeding **(Supplemental Figure 1)** Single cells were seeded at a density of 4000 cells/well in ultra-low attachment 24-well plates to induce mammosphere formation. Cells were cultured in normal culture media for 7 days and were subsequently counted, dissociated enzymatically in Accutase, (Stem Cell Technologies, Vancouver, Canada: #07922), and analyzed by flow cytometry. See flow cytometry methods for detailed information.

### Paclitaxel resistance studies

MCF10DCIS.com cells were treated with 100nM paclitaxel (Millipore Sigma, Burling, MA, USA #T7402-5MG) for 24 hours prior to analysis by flow cytometry. Dose was determined experimentally based on cell viability and cancer stem cell enrichment. Drug resistance was measured by label-retention and mammosphere formation as described above and %CD44+ (Biolegend: #103018) CD24- (Biolegend: #311104) of singlets was used as a molecular definition of drug-resistant stem cells. See label retention stem cell assay and flow cytometry methods for additional details.

### Flow cytometry

Single cell suspensions were generated from cultured cells via enzymatic harvest with Accutase. Mouse mammary tissues were minced with scalpels, digested in Click’s media containing 500 units/mL Collagenase II (Worthington, Columbus, OH, USA; LS004174) and Collagenase IV (Worthington, LS004186) and 20ug/mL DNAse (Worthington, LS002004) for 1 hour at 37°C with occasional trituration and strained through 70um filters to generate single cell suspensions. Single cells were stained at 4°C for 30 minutes and washed with PBS+2%FBS. Cells were filtered through 30um filters, diluted to <10^6^ cells/ml, and analyzed with the ZE5-YETI flow cytometer. Fixation/Permeabilization kit (BD Biosciences, Franklin Lakes, NJ, USA: #554715) was used for intracellular antigen staining. Data were analyzed with Kaluza software. Single stain, FMO, and unstained controls were used to set gates. Single cells were lineage depleted using CD45- (BD Biosciences: #564279)/CD31- (BD Biosciences: #745436) gating. Full gating strategy is included as Supplemental Figure 2.

### Immunohistochemistry

Tissues were formalin-fixed and paraffin-embedded as previously described(31). Hematoxylin and Eosin staining was used to define morphological features. Antigen retrieval was performed using target retrieval solution (Dako, Glostrup, Denmark; #S1699) for CC3(Cell Signaling Technologies, Danvers, MA, USA; #9661) and Perilipin(Cell Signaling: #3470) or (EDTA Dako cat# S2367) for pSTAT3 (Cell Signaling Technologies; #9145). ImmPRESS polymer anti-rabbit IgG secondary reagent (Vector Labs, San Francisco, CA, USA t#MP-7401-15) was used for secondary staining of CC3 and pSTAT3 stained tissues and anti-rabbit secondary (Dako #K4003) for perilipin stains. DAB (Vector Labs,; #SK-4105) and counterstaining were performed with hematoxylin (Vector Labs #H-3401). Alveolar area was measured on H&E-stained tissues in ImageJ on 10 alveoli/field; 5 fields/tissue. 10 fields/tissue were analyzed for positive CC3, pSTAT3, and perilipin stained tissues using CellSense Dimension software count and measure feature on regions of interest.

### Data Mining

Analysis of SEMA7A in breast cancer versus normal was performed on http://ualcan.path.uab.edu/index.html by selecting breast cancer samples from the The Cancer Genome Atlas (TCGA) (32, 33). 5-year relapse free survival curves for *Sema7a* (Affy ID: 230345_at) and *Itga6* (Affy ID: 201656_at) were generated in KM plot using default parameters and no restrictions. Patients split by median survival. For multi-gene analyses, genes were filtered by median expression(34).

### Statistical Analysis

Unpaired *t*-test and one-way ANOVA were performed in GraphPad Prism. Nonparametric equivalent analyses were used for samples with uneven distribution or unequal standard deviations. *p* values of less than 0.05 were considered significant. Error bars represent mean +/− standard error of the mean. All in vivo studies were replicated in at least 3 mice and in vitro studies performed in technical and biological triplicate with representative data presented.

## RESULTS

### Semaphorin-7a promotes anoikis resistance in normal MECs

To identify whether SEMA7A promotes survival of normal MECs we utilized two methods: transfection of FC-tagged SEMA7A into immortalized human (MCF10A) cells **(Supplemental Figure 3)** and addition of exogenous SEMA7A, which we purified from FC-tagged SEMA7A expressing cells, to wild-type MCF10A and MCF12A cells—another non-transformed MEC line. We found that genetic SEMA7A overexpression resulted in resistance to cell death, as measured by cleaved caspase 3/7, in a forced suspension assay, but not in attached conditions suggesting that SEMA7A specifically mediates resistance to cell death by anoikis(**Figure 1A)**. To determine the contribution of endogenous SEMA7A protein to anoikis resistance, we measured SEMA7A+ MECs by flow over time in attached and suspended conditions and found that SEMA7A+ MECs in attached conditions remain unchanged while SEMA7A+ cells become significantly enriched as early as 24 hours in detached conditions(**Figure 1B**). We also show that stimulation with exogenous purified SEMA7A protein in both MCF10A and MCF12A cells is sufficient to significantly decrease cell death and Inhibition of a known SEMA7A receptor, b1-integrin, significantly abrogated SEMA7A-mediated anoikis resistance **(Figure 1C and Supplemental Figures 4 and 5)**. We independently confirmed decreased cell death in detached MECs cultured with exogenous SEMA7A by flow cytometry for Annexin-V/7AAD (**Supplemental Figure 6**). We also examined pAKT in MECs cultured with SEMA7A using insulin as a positive control as the IGF-1 receptor (and insulin) can activate pAKT via b1-integrin in the mammary gland(35, 36). We observed percentages of pAkt expressing cells that were similar to those observed with insulin when cells were exposed to SEMA7A *in vitro* **(Figure 1C)**.

**Figure 1:**
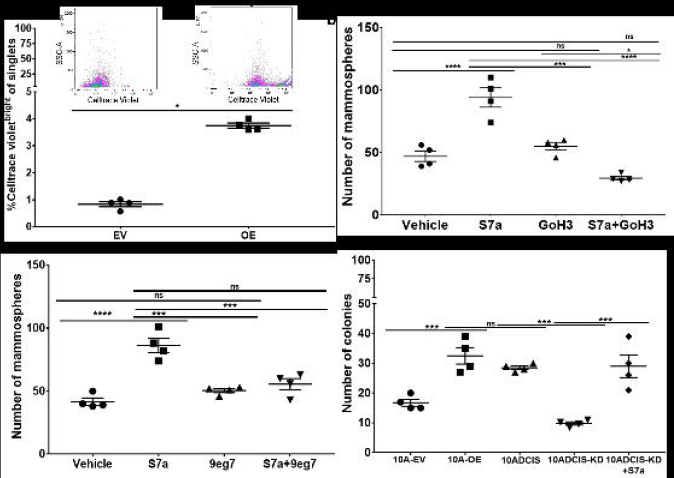
Semaphorin-7a promotes anoikis resistance in normal MECs. **a.** Cleaved caspase 3/7 (CC3/7) expression in MCF10A EV or S7a-OE cells after 24 hours in attached or forced suspension cultures. **b.** SEMA7A expression by flow cytometry in attached and forced suspension MCF12A cultures. **c.** CC3/7 in MCF12A cells in forced suspension +/− exogenous S7a protein and/or ITGB1 blocking antibody, 9eg7. **d**. Flow cytometry analysis of pAKT in insulin or S7a-treated MCF12A with representative flow plots. **e.** Flow cytometry for %SEMA7A+ mammary epithelial cells from tissues harvested from nulliparous (N) or recently weaned C57/BL6 mice at involution days 1-28 (I1-I28). **f.** Flow cytometry of live epithelial cells as measured by Aquazombie-EpCAM+ in mammary tissues harvested from recently weaned WT or Sema7a^tm1Alk/J^ mice. WT (I1: n=5, I2: n=6, I3: n=5, I6: n=4 glands/group) SEMA7A ^tm1Alk/J^ ( I1-I6: n=6 glands/group). Error bars indicate mean +/− standard error of the mean. Significance denoted as: p<0.05=*, p<0.01=**, p<0.001=***, p<0.0001=****.

To determine whether SEMA7A inhibits anoikis *in vivo*, we examined MECs harvested from age-matched nulliparous and force weaned WT animals for SEMA7A expression. We observed a low percentage of SEMA7A+EpCAM+ cells in the nulliparous mammary gland, which was enriched approximately 3-fold at involution day 3 when basement membrane degradation and extracellular matrix remodeling ensues. The percentage of SEMA7A+ epithelial cells decreased starting at involution day 4 and returned to pre-involution day 3 levels by involution day 6 **(Figure 1E)**. We also compared live epithelial cells by flow cytometry in age matched SEMA7A knockout mice (Sema7a^tm1Alk/J^) and WT C57Bl/6 mice at days 1, 2, 3, & 6 post-wean and observed a modest, but significant, reduction in MECs as measured by the percentage of Aquazombie-EpCAM+ cells **(Figure 1F**). To test whether the decrease in live cells during involution in Sema7a^tm1Alk/J^ animals was accompanied by changes to gland histology and programs normally activated during postpartum involution, we performed H&E and immunohistochemical analysis of whole mammary tissues isolated from Sema7a^tm1Alk/J^ dams and WT controls. We observed accelerated glandular regression, as evidenced by decreased alveolar area, increased pSTAT3, precocious appearance of cleaved caspase 3 (CC3) positive cells, and accelerated adipocyte reappearance in glands harvested from Sema7a^tm1Alk/J^ animals **(Figure 2 and Figure 3)** suggesting that loss of SEMA7A accelerates postpartum involution.

**Figure 2:**
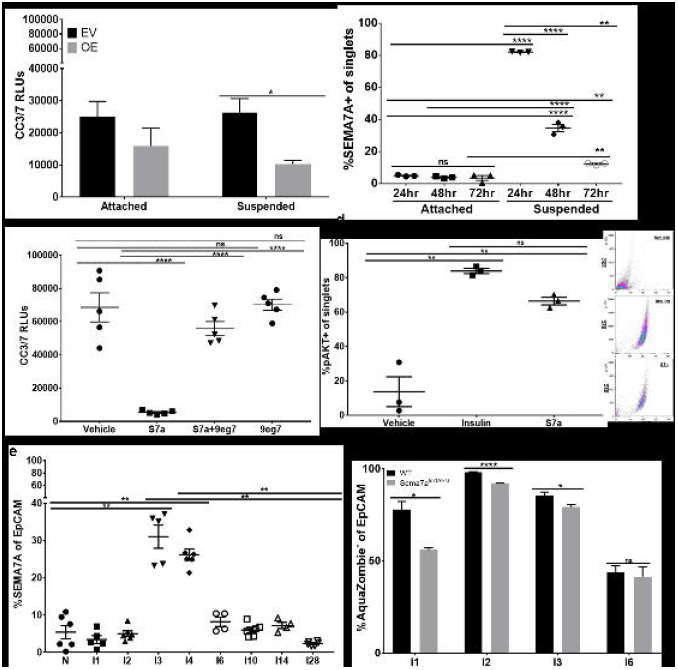
Loss of SEMA7A promotes histologic features of accelerated involution. Representative 20x images of #3 mammary glands from WT or SEMA7A ^tm1Alk/J^ mice at 1,2,3,&6 days of forced involution. **a.** Hematoxylin and Eosin, **b.** pSTAT3, **c.** cleaved caspase-3 and **d.** perilipin staining.

**Figure 3:**
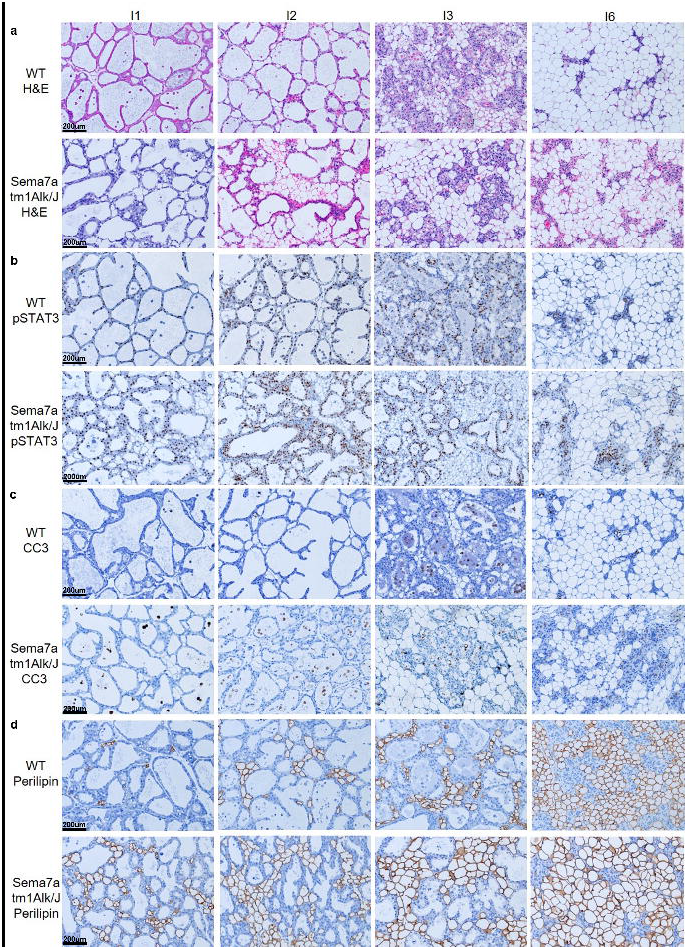
Loss of SEMA7A promotes alveolar collapse through alterations to cell survival and adipocyte repopulation. Immunohistochemical quantification of mammary glands from WT (I1: n=5, I2: n=6, I3: n=5, I6: n=4 glands/group) or SEMA7A ^tm1Alk/J^ ( I1-I6: n=6 glands/group) mice at 1,2,3,&6 days post forced involution. Quantification of **a.** alveolar area **b.** pSTAT3 **c.** cleaved caspase-3 and **d.** perilipin staining. Error bars indicate mean +/− standard error of the mean. Significance denoted as: p<0.05=*, p<0.01=**, p<0.001=***, p<0.0001=****.

### SEMA7A promotes anoikis-resistance via a6-integrin and activates AKT

MECs can be divided into four primary cell types: luminal, luminal progenitor (LPCs), basal, and basal progenitor cells (BPCs) and these populations can be resolved by flow cytometry based largely on their expression of epithelial cell adhesion molecule (EpCAM) and CD49f, also known as α6-integrin(37, 38). Since the magnitude of the changes observed in Figure 1E was relatively small, we used flow cytometry to resolve the four primary MEC populations to determine which cells are specifically affected by loss of SEMA7A during postpartum involution. We observed a significant decrease in EpCAM^hi^CD49f+ LPCs at all timepoints examined, which was not consistently observed in the other cell populations **(Figure 4A,B and Supplemental Figure 7)**. We also found that more than 80% of the live LPCs express SEMA7A at involution days 2-4 **(Figure 4C)**. Furthermore, SEMA7A+ cells are enriched in each MEC population at some point during postpartum involution except for mature luminal cells. **(Supplemental Figure 8)**. One factor that distinguishes the SEMA7A+ MEC populations from the SEMA7A-luminal cells is their expression of α6-integrin. β1-integrin is a known receptor that mediates SEMA7A cellular signaling(8), but the preferred alpha binding partner in MECs has not been identified. We quantified cells that co-express α6 and β1 in WT and SEMA7A^tmalk1/J^ animals and found a significant reduction during involution **(Figure 4D)**. Furthermore, we observed that over 90% of the α6 and β1 positive MECs express SEMA7A at the transition to the reversible phase of involution when basement membrane degradation and anoikis-induced cell death is initiated **(Figure 4E)**. To determine whether SEMA7A confers anoikis-resistance via α6-integrin dependent activation of pro-survival signaling, we cultured MCF12A cells with SEMA7A in the presence of the α6-integrin blocking antibody (GoH3) in forced suspension cultures. Consistent with our previous results, SEMA7A decreased CC3/7 activity, which was blocked by GoH3. SEMA7A treatment also resulted in enrichment for pAKT+ cells which was inhibited by α6 β1 blockade **(Figure 4F&G)**. Inhibition of both α6 and β1 did not increase CC3/activity or alter pAKT over vehicle alone in the absence of SEMA7A (**Figure 1C and 4G**).Taken together, our results suggest SEMA7A promotes anoikis-resistance during involution, in part, through α6β1and activation of pAKT.

**Figure 4:**
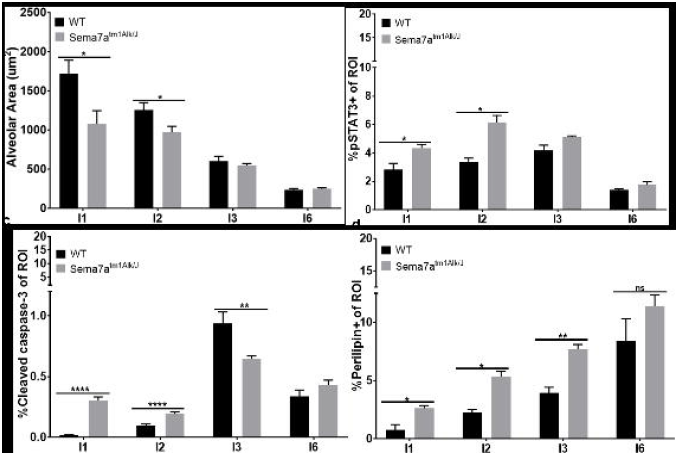
Semaphorin-7a promotes survival via integrins and enriches for pAKT+ cells. **a-e.** Flow cytometry analysis of mammary glands from WT (left, I1: n=4, I2: n=6, I3: n=5, I6: n=4 glands/group) and SEMA7A^tm1Alk/J^ (right, I1-I6: n=6 glands/group) mice. **a.** Stratification of MEC subtypes by EpCAM/CD49f staining. **b.** %EpCAMhiCD49f+ LPCs of EpCAM+ cells. **c.** %SEMA7A+ of live (Aquazombie-) EpCAMhiCD49f+ LPCs. **d.** %ITGB1+ITGA6+ of EpCAM cells. **e.** %SEMA7A of ITGB1+ITGA6+ cells. **f.** CC3/7 expression in MCF12A cells +/− S7a and/or GoH3. **g.** %pAKT in MCF12A cells +/− S7a and/or 9eg7/GoH3, n=3. Error bars indicate mean +/− standard error of the mean. Significance denoted as: p<0.05=*, p<0.01=**, p<0.001=***, p<0.0001=****.

### SEMA7A promotes stem cell phenotypes, cellular transformation, and chemoresistance

Anoikis-resistance is rarely observed in normal epithelial cells; rather, it is a characteristic of stem cells and tumor cells. Given our findings that SEMA7A promotes anoikis-resistance in normal MECs, we tested whether it also promotes functional features of mammary stem cells using cultured MECs. Previously identified functional properties of mammary stem cells include a quiescent or slowly proliferative phenotype and their ability to survive and proliferate in anchorage-independent conditions where they give rise to clonal spheroids. These spheroids, also known as “mammospheres”, allow for identification of stem cells as they only undergo one or two rounds of division before reentering quiescence, allowing them to be identified based on their ability to retain fluorescent dyes that are normally progressively lost by dilution in the actively proliferating cells(39–41). We used a cell trace violet dye and mammosphere assay to identify functional mammary stem cell phenotypes *in vitro* and found that SEMA7A expressing MCF10A cells are enriched for a quiescent, or senescent, cell population **(Figure 5A)**. We also measured mammosphere formation as a readout for self-renewal in cells cultured with exogenous SEMA7A in the presence or absence of function blocking antibodies for α6 and β1 and found that increased sphere formation occurred with SEMA7A which was abrogated with the inhibitors **(Figure 5B&C)**. Secondary sphere formation was also increased in MCF10A and MCF12A cells cultured with SEMA7A **(Supplemental Figure 9)**. We then tested the ability of SEMA7A to promote cell growth in anchorage-independent conditions using a soft agar assay— a classical measurement of cellular transformation. We found that overexpression of SEMA7A in MCF10A cells is sufficient to increase growth in soft agar to levels similar to a Ras-transformed variant of these cells, MCF10DCIS.com.

**Figure 5:**
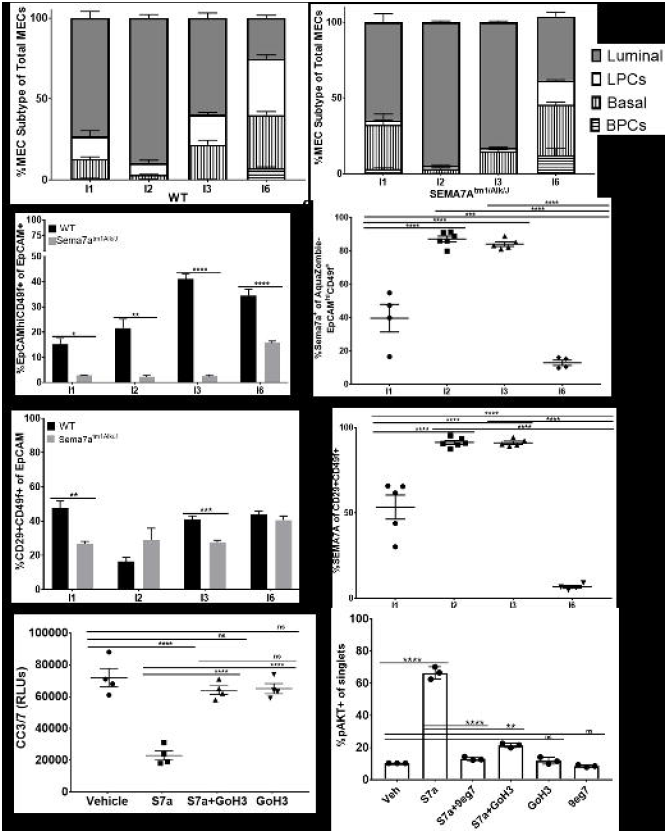
SEMA7A and ITGB1/ITGA6 promote features of transformation and chemoresistance. **a**. %Celltrace violet bright of singlets in MCF10A EV and OE cells. **b,c.** Number of mammospheres formed by MCF12A cells +/− S7a and GoH3 or 9eg7. **d.** Number of colonies formed in soft agar assay.

To determine a role for SEMA7A in breast cancer, we examined *SEMA7A* expression in breast cancers in The Cancer Genome Atlas (TCGA) to show that *SEMA7A* is significantly upregulated in primary tumors **(Figure 6A)**, is higher in all stages **(Figure 6B)**, is upregulated in breast cancers in women of all ages **(Figure 6C)** and is expressed in every major subtype of breast cancer with highest expression occurring in the triple-negative class when compared to normal breast tissues **(Figure 6D and Supplemental Figure 10)**. We also utilied Kaplan-Meier analysis to examine relapse free survival (RFS) to determine whether coordination between *SEMA7A* and *α6-integrin* impacts patient recurrence. We observed that expression of each molecule individually did not decrease RFS, but co-expression of *SEMA7A* in *α6*^high^ tumors did (HR=1.5; 95% CI=1.18-1.9; p=0.00075) **(Figure 6F and Supplemental Figure 11)**. Taken together, these data suggest a clinically relevant mechanism by which SEMA7A and a6-integrin may promote cellular phenotypes associated with tumor progression and chemoresistance.

**Figure 6:**
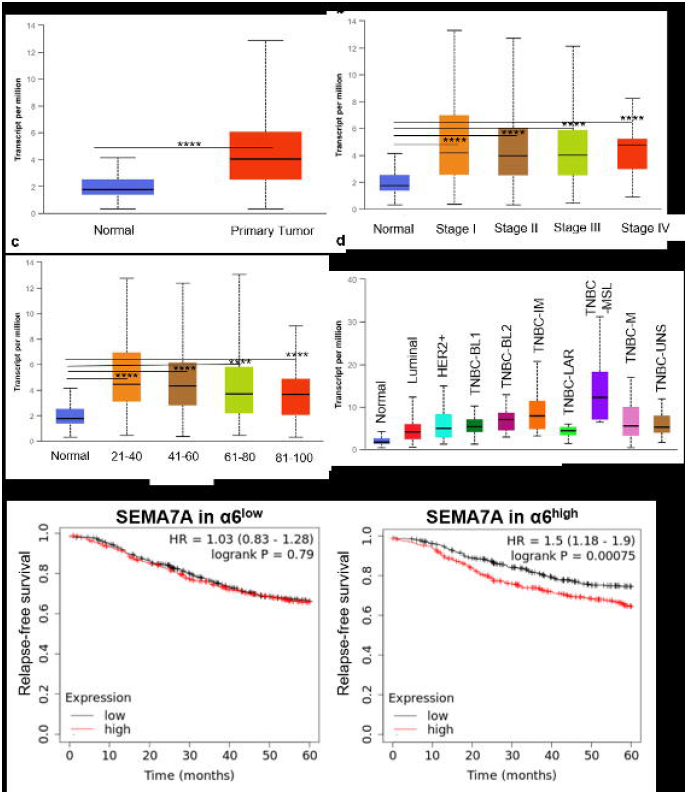
*Sema7a* is upregulated in multiple breast cancers and predicts for decreased relapse-free survival when expressed with a6-integrin. **a-d.** TCGA data for *Sema7a* in a. normal (n=114) vs primary tumor (1097) **b.** normal versus stages I-IV (I: n=183, II: n=615, III: n=247, IV: n=20) **c.** normal versus cancer stratified by age (21-40: n=97, 41-60: n=505, 61-80: n=431, 81-100: n=54) , **d.** normal versus breast cancer subclass (luminal: n=566, HER2+: n=37, TNBC-BL1: n=13, TNBC-BL2: n=11, TNBC-IM: n=20, TNBC-LAR: n=8, TNBC-MSL: n=8, TNBC-M: n=29, TNBC-UNS: n=27) Error bars indicate ??? Significance denoted as: p<0.05=*, p<0.01=**, p<0.001=***, p<0.0001=****. **e.** Kaplan-Meier 5-year relapse-free survival analysis for *Sema7a in Itga6* low (n=2032) or high (n=2032) expressing patients.

## DISCUSSION

Despite published roles for SEMA7A in breast cancer progression, the current study is, to our knowledge, the first investigation into the role of SEMA7A in normal MEC survival during postpartum mammary gland involution. In 2004, Clarkson and Watson published a comprehensive analysis of gene expression across mammary gland development in which they identified roles for both death receptors and immunomodulation during specific phases of postpartum development including the first and second phases of involution(43). In this highly impactful study, they identified seven basic profiles of gene expression across the lactation-involution cycle. *SEMA7A* was identified in the Inv1 cluster, which was upregulated at the transition between lactation and involution. In this study, we assessed cell surface SEMA7A protein expression by flow cytometry in recently weaned mice to show its expression during the initial phases of postpartum involution. We found that the proportion of MECs expressing SEMA7A peaks at involution day 3, marking the transition from the reversible to the irreversible phase of mammary gland remodeling. During the irreversible phase, MECs lose contact with the basement membrane, primarily composed of laminin, and die by anoikis. Since anoikis is one of the hallmarks of the irreversible phase of mammary involution, and our lab has published that SEMA7A overexpression in multiple breast cancer cell lines decreases anoikis and, most recently, that SEMA7A expression increases over time when cells are detached from matrix, we predicted that expression of SEMA7A would promote anoikis resistance during the second phase of irreversible involution(13, 44). We now also show that treating cells with exogenous SEMA7A can confer resistance to anoikis in cultured human MECs suggesting that SEMA7A may signal to neighboring cells to promote resistance to cell death. This is consistent with recent data showing that, in human breast tissues, entire acini are lost during involution rather than a percentage of cells from each acini(45). We propose that SEMA7A promotes anoikis resistance, in part, by signaling to detached cells in a manner similar to matrix molecules, thereby mimicking attachment. Consistent with this hypothesis, we show that exogenous SEMA7A increases pAKT, and cell survival, in cultured MECs in a manner that is dependent, in part, on α6 and β1-integrin, which is normally the integrin receptor pair for pro-survival signaling via ligation by laminin in the mammary gland(6, 24). While β1-integrin is a known mediator of SEMA7A signaling in a variety of contexts, few studies have addressed the alpha integrin that cooperates with β1 to mediate SEMA7A-activated signaling. Our results presented herein suggest, for the first time, that SEMA7A may mimic the effects of laminin to promote anoikis resistance via α6β1 heterodimers in MECs. One limitation of our study is that we have not directly assessed the effect of SEMA7A on anoikis-resistance in the presence of an AKT inhibitor which would directly establish a relationship between SEMA7A and AKT-dependent anoikis-resistance. However, literature suggests that AKT is important for promoting anoikis resistance, via inhibition of BIM, and that loss of AKT increases cell death by anoikis. Therefore, the ability of SEMA7A to enrich for pAKT+ cells suggests it may be sufficient to induce anoikis-resistance through published AKT-dependent mechanisms, but that SEMA7A is not the only molecule that can activate AKT to promote cell survival(46–48). Additional investigation is required to further understand the role of SEMA7A in this pathway.

Anoikis resistance is a hallmark of stem cells and breast cancer cells, particularly those that result in lethal metastasis as these cells must survive in the absence of extracellular matrix signaling as they travel to and throughout the vasculature to seed distant tumors(49). Since tumor cells are known to hijack normal developmental programs, we postulated that sustained expression of SEMA7A in MECs, particularly in postpartum women, could result in pro-tumor characteristics that are similar to those induced by Ras—a known oncogene. This hypothesis was based on our recent publication showing that SEMA7A expression is higher in normal adjacent breast tissue from postpartum women and our identification of a cut-point of expression that predicts for recurrence in women with PPBC(50). Our hypothesis is supported by our findings that SEMA7A mediates hallmarks of cellular transformation and stemness in non-transformed cultured MECs. While others have shown that SEMA7A is required for Ras dependent induction of EMT(51), we now show that growth in soft agar in Ras transformed variants of MCF10A cells is dependent on SEMA7A and that SEMA7A can promote growth of non-transformed cells in soft agar(26, 27, 30). Therefore, we predict that SEMA7A promotes cellular transformation and tumor initiation in breast cancer patients and, in particular, in PPBC patients(22, 30). A question that remains unanswered is whether sustained expression of SEMA7A is causative for malignancy or whether SEMA7A expression in the tumor influences normal adjacent cells to express SEMA7A thereby promoting tumor progression and metastasis. Evidence for the latter comes from our studies showing that SEMA7A can affect multiple aspects of the tumor microenvironment including collagen and fibronectin deposition, lymphangiogenesis and macrophage infiltration, as well as neighboring cells such as fibroblasts(26, 30). Additionally, we provide evidence that SEMA7A expressing cells upregulate drug resistant cell-surface markers(23) and therefore may not respond to conventional chemotherapy; this, along with our analysis of RFS in a large patient dataset revealing that co-expression of *ITGA6* and *SEMA7A* significantly influences probability of relapse suggests that SEMA7A can not only initiate tumor formation, but also influence clinical outcomes. Future studies will investigate these important questions to identify how to best prevent and treat PPBC and all patients with α6-integrin+/SEMA7A+ breast cancers.

Finally, to date, the majority of research into PPBC has focused on how the tissue microenvironment in the postpartum breast drives metastasis with lesser focus on cellular mechanisms at play in the MECs that may also support transformation(50). As more than half of all breast cancers in patients under 40 can be defined as PPBC, which are highly metastatic, understanding the mechanisms at play in postpartum breast tissue is, thus, critically important for identifying molecular drivers of PPBC. In this study, we have identified a known tumor-promotional protein, SEMA7A, as being temporally expressed during the irreversible phase of postpartum mammary gland involution where it promotes anoikis resistance ,and we have shown for the first time that SEMA7A can promote properties of transformation, stemness, and metastasis in non-cancerous cells.

## ACKNOWLEDGEMENTS

This work was supported by NIH/NCI R01 CA211696-01A1 (to T.R. Lyons), NIH/NCI R01CA211696-02S1(to Taylor Rutherford), ACS RSG-16-171-010CSM (to T.R. Lyons), and Department of Medicine Outstanding Early Career Scholars Award (to T.R. Lyons). The authors acknowledge the University of Colorado Cancer Center Flow Cytometry Core for use of equipment and technical assistance and the University of Colorado Cancer Center Support Grant (P30CA046934). The authors also thank Alexander Stoller for assistance with animal studies and Veronica Wessels for histological expertise. Contents are the authors’ sole responsibility and do not necessarily represent official NIH views.

The authors declare no potential conflicts of interest.

